# Live Cell Partial Wave Spectroscopic microscopy: Label-free Imaging of the Native, Living Cellular Nano-architecture

**DOI:** 10.1101/061747

**Authors:** L. M. Almassalha, G. M. Bauer, J. Chandler, S. Gladstein, L. Cherkezya, Y. Stypula-Cyrus, S. Weinberg, D. Zhang, P. Thusgaard Ruhoff, H. Roy, H. Subramanian, N. Chandel, I. Szleifer, V. Backman

**Affiliations:** Department of Biomedical Engineering, Northwestern University, Evanston, Illinois, 60208, USA; Department of Medicine, Northwestern University Feinberg School of Medicine, Chicago, Illinois, 60611, USA; Institute of Technology and Innovation, University of Southern Denmark, Campusvej 55, DK-5230 Odense M, Denmark; Section of Gastroenterology, Boston Medical Center/Boston University School of Medicine, Boston, Massachusetts, 02118, USA; Chemistry of Life Processes Institute, Northwestern University, Evanston, Illinois, 60208, USA

**Keywords:** Live Cell Imaging, Chromatin structure, DNA damage, Cellular Metabolism

## Abstract

The organization of chromatin is a regulator of molecular processes including transcription, replication, and DNA repair. The structures within chromatin that regulate these processes span from the nucleosomal (10nm) to the chromosomal (>200nm) levels, with little known about the dynamics of chromatin structure between these scales due to a lack of quantitative imaging technique in live cells. Previous work using Partial Wave Spectroscopic (PWS) microscopy, a quantitative imaging technique with sensitivity to macromolecular organization between 20-200nm, has shown that transformation of chromatin at these length scales is a fundamental event during carcinogenesis. As the dynamics of chromatin likely play a critical regulatory role in cellular function, it is critical to develop live-cell imaging techniques that can probe the real-time temporal behavior of the chromatin nano-architecture. Therefore, we developed a live cell PWS technique which allows high-throughput, label-free study of the causal relationship between nanoscale organization and molecular function in real-time. In this work, we employ live cell PWS to study the change in chromatin structure due to DNA damage and expand on the link between metabolic function and the structure of higher-order chromatin. In particular, we studied the temporal changes to chromatin during UV light exposure, show that live cell DNA binding dyes induce damage to chromatin within seconds, and demonstrate a direct link between higher-order chromatin structure and mitochondrial membrane potential. Since biological function is tightly paired with structure, live cell PWS is a powerful tool to study the nanoscale structure-function relationship in live cells.

**Significance Statement:** Chromatin is one of the most critical structures within the cell because it houses most genetic information. Its structure is well understood at the nucleosomal (<20nm) and chromosomal (>200nm) levels, however, due to the lack of quantitative imaging modalities to study this organization, little is known about the higher-order structure between these length scales in live cells. We present a label-free technique, live cell Partial Wave Spectroscopic (PWS) microscopy with sensitivity to structures between 20-200nm that can quantify the nano-architecture in live cells. With this technique, we can detect DNA fragmentation and expand on the link between metabolic function and higher-order chromatin structure. Live cell PWS allows high-throughput, label-free study of the causal relationship between nanoscale organization and molecular function in live cells.

## Introduction

Every cellular and extracellular structure has a complex nanoscale organization ranging from individual macromolecules that are a few nanometers in size (e.g. protein, DNA) to macromolecular assemblies that are tens to hundreds of nanometers in size (e.g. cell membranes, higher-order chromatin structure, cytoskeleton, and extracellular matrix fibers). A major scientific challenge is to understand these macromolecular structures, specifically their function and interactions in structurally and dynamically complex living cellular systems. To meet these goals, the ideal live cell imaging technology would satisfy five key requirements: (1) nanoscale sensitivity (<200nm), (2) label-free (3) non-perturbing (4) quantitative, (5) high-throughput, and (6) molecularly informative.

Current approaches are unable to meet all these criteria alone. The breakthroughs in super-resolution fluorescence microscopy (SRM) have enabled new imaging technologies capable of providing unprecedented molecular identification at the highest resolutions currently available in live cells, but require the use of exogenous fluorophores to visualize macromolecular structures (1–3). For some applications, these labels are indispensable to achieve molecular specificity. However, there are significant drawbacks to the exclusive use of molecular labels for studies of cellular structure and function. Exclusively label-based SRM approaches are limited by the number of possible imaging channels, the high label-densities required, the high light intensities utilized during imaging, and by the introduction of possible artifacts due to the labels themselves, especially at the high densities required for nanoscale resolution (4, 5). In the study of macromolecular organization, current imaging approaches have significant drawbacks as macromolecular structures are often composed of dozens to hundreds of distinct molecules and often include different subtypes such as lipids, proteins, nucleic acids, and carbohydrates, some of which are difficult to directly label. Due to these limitations, phase contrast microscopy is still the most widely used label-free imaging modality for live cell experiments. While this technique is fast and improves contrast to visualize live cells, its diffraction-limited resolution cannot provide any insights into the macromolecular nano-architecture. As such, currently no label-free optical technique exists to measure the nano-architectural properties of live cells below the diffraction limit.

One prominent area of biological research with a demonstrated need for label-free, nanoscale sensitive imaging is the investigation of the structure-function relationship of chromatin. Chromatin organization (which is comprised of DNA, histones, and hundreds of other conjugated proteins and small molecules such as RNA) involves a hierarchy of length scales ranging from 10nm in nucleosomes to hundreds of nanometers for chromosomal territories (6, 7). The physical nanostructure of chromatin is regulated by numerous molecular factors, including the primary DNA sequence composition, differential methylation patterns, histone modifications, polycomb and cohesin protein complexes, RNA and DNA polymerases, long non-coding RNA, etc, and non-molecular factors, such as crowding, ionic conditions, and pH. Due to this complexity and the lack of existing optical techniques that can rapidly sample information below 200nm, little is known about the higher-order chromatin structure between these length scales or their dynamics in live cells. Results from fixed cell imaging techniques, such as electron microscopy or SRM, have shown that chromatin between 20-200nm is first organized into poly-nucleosomal 10 nanometer fibers, and in certain conditions, these fibers have been shown to assemble into 30nm clusters (8–10), although the existence of the 30nm fiber is a subject of an active debate. At length scales between 100-200nm, recent work using SRM has shown a power-law relation in the organization of chromatin, with domains of highly-dense, inactive chromatin localizing within a few-hundred nanometers of transcriptionally active sites (11). In conjunction, molecular techniques such as chromosomal capture methods (3C and Hi-C) have shown that the higher-order organization of chromatin above single nucleosomes and below the structure of chromosomal territories follows this same power-law fractal organization. These methods have shown that topology of this higher order organization is correlated with the regulation of gene transcription (12–14) and capable of evolving rapidly under stress conditions to globally regulate the expression of genes (15). Critically, these observed changes in chromatin structure have recently been linked to the regulation of genes often implicated in oncogenesis(16).

In cancer, it is increasingly clear that changes in chromatin topology at all length scales are a critical determinant of tumor formation, aggressiveness, and chemoresistance. One of the primary features of tumorigenesis is a shift in the fractal physical organization of chromatin, correlating both with the formation of tumors and their invasiveness. In early carcinogenesis, we have previously applied a fixed cell imaging technique, Partial Wave Spectroscopic (PWS) microscopy and transmission electron microscopy, to detect physical changes in the chromatin nano-architecture, indicating that the topology of chromatin is a critical component in cellular function at the earliest stages of tumor formation (17). PWS microscopy allows examination of the intracellular organization concealed by the diffraction limit with length scale sensitivity in the range of 20-200nm, the range at which existing label-free live cell imaging techniques are deficient, due to the relationship between the nanoscale spatial variations of macromolecular density and the resulting variance in the spectrum of backscattered light (18, 19). The novelty is grounded in a previously overlooked difference between resolution and detectability. While sub-diffractional structures are not *resolved* by PWS, they are still *detected* by analyzing the spectrum of elastically scattered light to provide quantitative contrast.

In this work, we set out to create a label-free live cell microscopy method based on interference principles used in PWS cytology, thereby creating a tool to directly study the dynamic nanoscale topology of live cells, with a specific focus on studying real-time changes in chromatin organization. We sought to develop an ideal extension of PWS in live cells that would (1) provide *nanoscale sensitivity* to structures between 20-200nm, (2) use *label-free contrast* to capture nanoscopic information, (3) be *non-perturbing to biological samples* by using low power illumination and label-free contrast, (4) *quantify* the cellular nanoarchitecture, and (5) rapidly capture the temporal evolution of nanoscale structures, providing contrast in multiple cells in seconds. For studies aimed at understanding the overall structure-function relationship in eukaryotic cells, employing SRM approaches alone would not be suitable. The power of live cell PWS is its unique ability to work in conjunction with existing label-based technologies to provide the structural context for molecular interactions, thereby greatly expanding our understanding of the molecular behavior in live cells (20). With this approach, we demonstrate the ability to measure the nano-architecture in live cells in seconds. Specifically, we explore changes to the cellular nano-architecture due to UV light exposure, show that live cell DNA binding dyes transform chromatin within seconds, and demonstrate a direct link between higher-order chromatin structure and mitochondrial membrane potential.

## Results and Discussion

The live cell PWS instrument is built into a commercial inverted microscope equipped with a broadband illumination and a tunable spectral collection filter [**Supplemental Figure 1**]. With this configuration, the live cell instrument utilizes the glass-cell interface to produce the requisite interference signal that allows for the study of underlying nanoscale structure. In brief, the spatial fluctuations of refractive index (RI) produced by the macromolecular density distribution cause backscattering of incident light waves from the sample. Optical interference of the back-propagating light results in wavelength-dependent fluctuations in the acquired spectrally-resolved microscope image. The standard deviation of these spectra (Σ) quantifies the internal structure of the sample with nanometer sensitivity (18, 19). In cells, there are numerous variations in macromolecular density due to the spatial organization of macromolecules. Quantification of this nano-molecular density distribution is given by the statistical parameter, Σ, at each diffraction-limited pixel (18, 19).

Σ, and the Disorder Strength (*L*_*d*_ which is Σ normalized by sample thickness), are proportional to two crucial characteristics of molecular organization at deeply subdiffractional length-scales: the characteristic length scale of macromolecular organization (*L*_*c*_), and the standard deviation (δn) of the density (18, 19). In a fractal media, such as chromatin, the characteristic length scale of macromolecules can be alternatively evaluated through the fractal dimension, D, which is proportional to Σ. Thus, Σ measured from chromatin senses nanoscale changes in its fractal organization. Critically, increase in heterogeneity (i.e. differential compaction) within chromatin by definition produces an increase in D, δn, or *L*_*c*_. This relation is derived from the properties of fractal media with conserved mass and volume – as compaction increases locally, the variations in mass density (heterogeneity) must also increase. Previous molecular dynamics simulations have further confirmed that increases in δn\**L*_*c*_ correspond to an increase in macromolecular compaction, and experimental results have shown that this increase within the nucleus quantitatively describes an increase in chromatin heterogeneity (17, 21, 22).

As a representation of the nanoscopic topology detected by live cell PWS, we used as a model 10nm “beads on a string” chromatin fibers (**Figure 1a&b**) as has been previously described by Kim *et al* (22). In this model, we consider changes in the nanoscopic structure of higher-order chromatin that have the same nanoscopic average mass density but have starkly different nanoscale organization: differentially compacted (**Figure 1a**) and diffusely compacted (**Figure 1b**) DNA fibers. In both cases, images produced from conventional light microscopy techniques cannot capture information about the nanoscale topology these differential states (**Figure 1c&d**). Likewise, while PWS cannot resolve the structures directly, it provides information about their sub-diffractional organization. To demonstrate sensitivity to these structures, E is computed directly as described by the work of Cherkeyzyan *et. al* accounting for the physical properties of the live cell system (18, 19). As is shown in **Figure 1e**, differentially compacted chromatin (**Figure 1a**) produces a much higher Σ than diffusely compacted chromatin (**Figure 1f**). Consequently, regions that result in high Σ in live cells would be the heterogeneous, differentially compacted regions likely resulting from the formation of local heterochromatin domains neighboring decompacted euchromatin (**Figure 1a**). Conversely, homogeneous regions of chromatin would result in low Σ.

**Figure 1:**
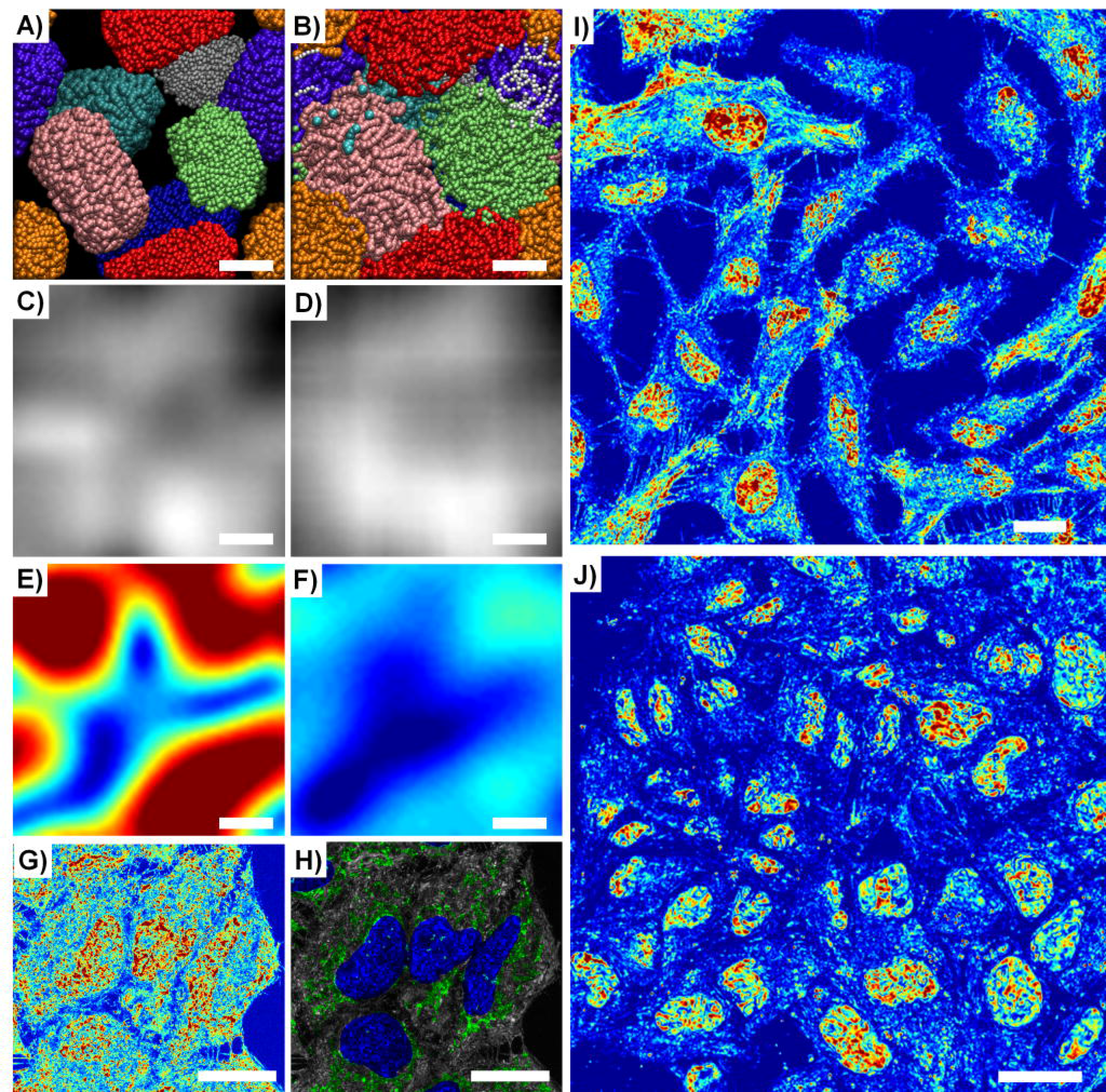
Live cell PWS rapidly provides quantitative nanoscale structural information of living cells. A&B) Orthographic z-axis projection of molecular dynamics simulations of chromatin as a 10nm “beads on a string” polymer capturing (**A**) differentially compacted (*l*_*c*_ = 70nm) and (**B**) diffusely compacted chromatin (*l*_*c*_ = 20nm). Scale bar represents 100nm. **C&D**) Calculated transmission microscope image captured by (**C**) conventional bright-field microscope from differentially compacted chromatin in (**A**) and (**D**) of diffusely compacted chromatin in (**B**). Images were produced by calculating the average mass density at each pixel and a Gaussian PSF of 250nm was applied to simulate a conventional microscope. Grid size of the simulations was 10nm. **E&F**) Calculation of Σ captured by live cell PWS from differentially compacted chromatin in (**A**) and diffusely compacted chromatin in (**B**). Σ was calculated directly from the distribution of mass within configurations shown in **A&B** with Σ=0.01-0.065. **G**) Representative pseudo-colored Live cell PWS image of HeLa cells with 63x oil immersion lens, NA=1.4 with Σ scaled to range between 0.0125 to 0.065. **H**) Co-localization of fluorescence with Live cell PWS image showing mitochondria (green), nuclei (dark blue), and mitochondria-nucleus overlap (light blue). Scale bar is 20μm. **J&K**) Representative pseudo-colored Live cell PWS image of (**J**) HeLa cells and (**K**) MES-SA cells demonstrating the capacity to capture nanoscopic information from dozens of nuclei in seconds with Σ scaled to range between 0.01 to 0.05 in J and 0.01 to 0.065 in K.

While this new instrument configuration was optimized to allow live cell imaging with multi-modal acquisition, including wide-field fluorescence and phase contrast microscopy, it has an appreciably weaker reference-interference signal than that produced in traditional PWS cytology and a much higher objective collection numerical aperture. Therefore, we validated the nanoscale sensitivity of live-cell PWS by using rigorous Finite-Difference Time-Domain computations [**Supplemental Figure 2**] to numerically solve Maxwell’s equations without approximations simulating the nanoscale-complex spatial distribution of molecular density in live cells. These computations were employed to study the effect of the RI mismatch using sapphire as a high-RI substrate on the interference signal [**Supplemental Origin of Interference Spectrum**], and to compare the effect of numerical aperture on Σ [**Supplemental Figure 3**]. The FDTD simulations enabled us to optimize the configuration of signal acquisition in order to provide nanoscale sensitivity to intracellular structure at length scale between 20 and 200 nm.

Without the use of exogenous labels, we can achieve high-contrast images using Σ that delineate nuclei from cytoplasm due to the intrinsic differences in their nano-architecture [**Figure 1g and Supplemental Figure 4** for additional comparisons in multiple cell lines, including primary cell lines]. Due to its multimodal design, exogenous and endogenous labels can be subsequently used to co-localize specific molecular markers or organelles [**Figure 1h**]. Live cell PWS acquisition yields a three-dimensional data cube, I(λ,x,y), where λ is the wavelength and (x,y) correspond to pixel positions across a 10,000μm^2^ field of view, allowing multiple cells to be imaged simultaneously. Acquisition of the full cell-reference interference spectrum (500-700nm) for spectral analysis takes under 30s, with each wavelength collection produced from < 100ms exposures. Using a reduced wavelength approach to sub-sample the interference spectrum, this can be further reduced to under 2s per acquisition (23). Even with full spectral collection, as demonstrated in **Figure 1j&k** (and **Supplemental Figure 4**), live cell PWS provides rapid, quantitative visualization of cellular structures within a single field of view for dozens of cells simultaneously for multiple cell lines (**Figure 1j**, 20 HeLa cells captured in ~30 seconds and **Figure 1k**, 36 MES-SA cells captured in ~15 seconds). Indeed, one of the most critical features of this rapid acquisition is the capacity to directly study the underlying heterogeneity of both chromatin structure and its temporal evolution within the cell population over time. Likewise, as a label-free technique using low illumination intensity, live cell PWS allows for the study of various time evolving processes on the structure of cells in general, and chromatin in particular for different cell types over extended periods of time.

Live cell PWS has a broad utility as a tool for studying the complex relationships between cell function and chromatin nano-organization. As the initial demonstration, we explored live-cell PWS to rapidly quantify the changes in the nanoscale chromatin structure due to DNA damage. As a demonstration of its ability to detect rapid changes specific to chromatin (within seconds), we explored the transformation of the higher-order chromatin structure secondary to DNA fragmentation using the DNA binding dye, Hoechst 33342. Damage to DNA results in the formation of DNA fragments and double stranded breaks (DSBs)(24–26). This damage, in turn, leads to apoptosis or mutagenic transformation. In cancer therapy, many drugs eliminate tumor cells by causing an unbearable accumulation of DNA damage and/or by activating the intrinsic apoptotic pathways (27, 28). Therefore the identification of DNA fragmentation and understanding of the time evolution of chromatin structure in response to damage is crucial to both understanding DNA repair mechanisms and to identifying chemotherapeutic efficacy.

Current techniques to study these processes require cell fixation, such as immunofluorescent identification of the rapidly phosphorylated histone H2A.X (γ-H2A.X) subunit (26) or transfection using photoactivatable proteins (29, 30). Furthermore, fluorescent visualization of cell viability for drug screening often requires the use of cell permeant minor-groove binding dyes. However, it has been reported that these minor-groove DNA binding dyes, including Hoechst 33342, induce DNA fragmentation by disrupting the activity of topoisomerase I(31, 32). These effects are observed independently from the fluorescence excitation of the dye, but are accelerated upon UV excitation(32). Consequently, no methods currently exist with the capability for the real-time study of changes to chromatin higher-order structure due to DNA damage or the overall dynamics of the damage response in unlabeled live cells.

Using live cell PWS, we show for the first time that the addition of Hoechst 33342 to living mammalian cells transforms the nano-organization of chromatin at the time scale required for imaging and that subsequent excitation induces fragmentation of the chromatin nano-architecture within seconds. This is apparent, as we observe an overall decrease in the Σ after irradiation, suggesting homogenization and decompaction of chromatin across the entire nucleus [**Figure 2a**]. Additionally, these effects persist for longer durations, lasting at least 15 minutes indicating that the once fragmented, chromatin in the presence of the dye does not immediately reassemble suggesting these changes could be irreversible. To control for the effects of ionizing UV radiation required for Hoechst excitation, we performed a mock-staining (M-S) experiment where we compared the nuclear changes in cells incubated with Hoechst 33342 to those exposed to UV light alone. In the M-S cells, there was not an observable change in cellular or chromatin structure during the short illumination time required for Hoechst excitation, indicating preservation of the original chromatin structure [**Figure 2b&c**]. Quantitatively, M-S cells showed no significant change in mean-nuclear E after a few seconds of UV exposure, whereas the Hoechst-stained cells display a 17.01% decrease in HeLa (99% confidence interval Hoechst (-18.5%, −15.6%), p-value < 0.001) between mock and Hoechst stained cells with n = 146 cells from 11 independent experiments for Hoechst stained cells and n = 68 cells from 6 independent experiments for M-S cells [**Figure 2b&c**]. In Hoechst-stained cells, all nuclei demonstrate a negative change in the mean nuclear Σ after UV exposure, whereas the M-S cells display a narrow, zero centered distribution after UV exposure [**Figure 2e**]. In both M-S and Hoechst-stained cells, cytoplasmic Σ did not change following UV exposure (p-value > 0.05). Similar results were observed for Chinese Ovarian Hamster (CHO) cells with M-S cells displaying no change, whereas Hoechst stain cells experience a −7.1% decrease (99% confidence interval Hoechst (-9%, −5%), p-value <0.001 between M-S and Hoechst stained cells, n=127 cells for M-S, n=87 for Hoechst-stained from 5 independent experiments each), demonstrating this effect occurs independent of the cell type [**Supplemental Figure 3**].

**Figure 2:**
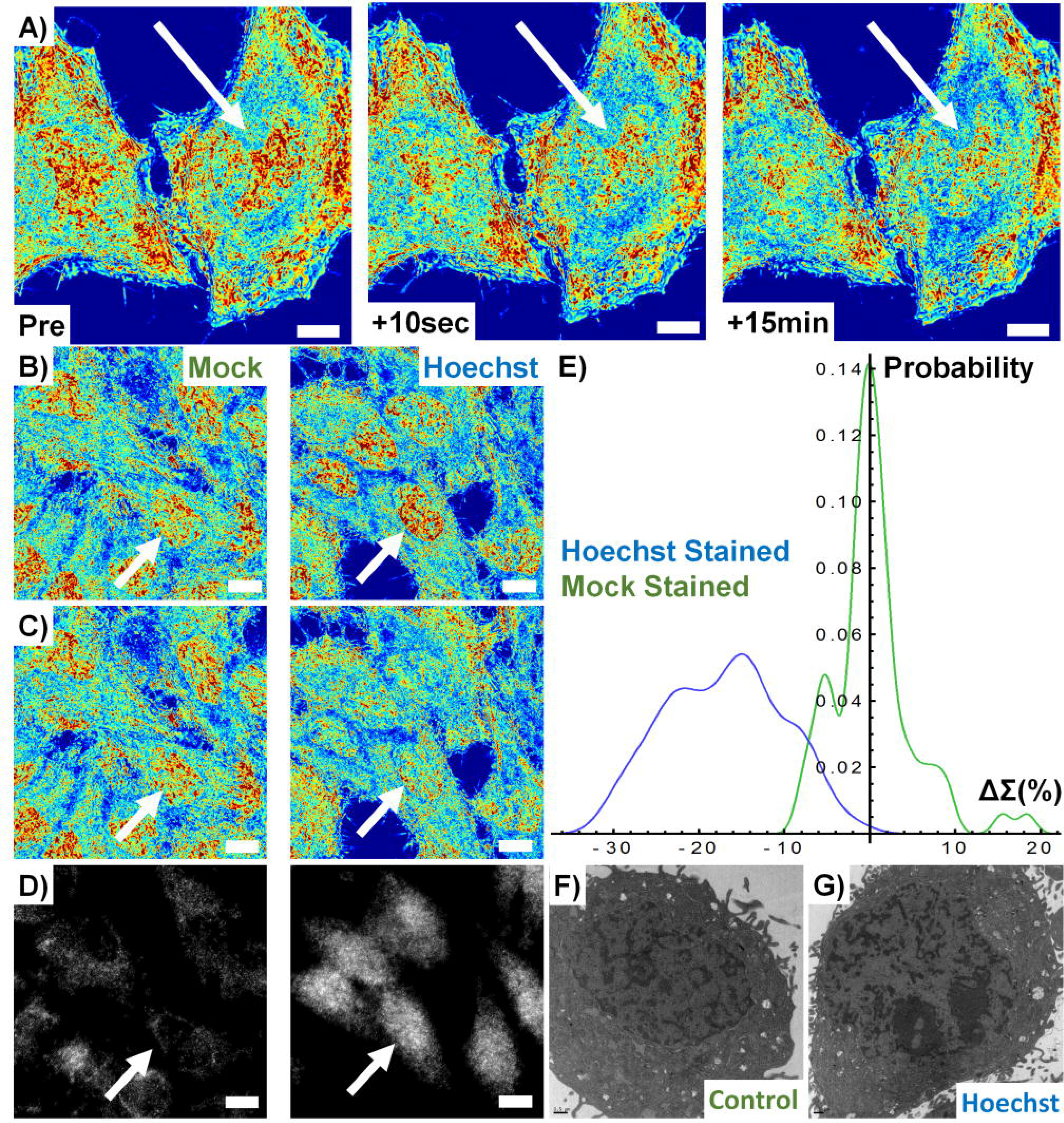
Hoechst excitation induces rapid transformation of chromatin nano-architecture. Row A) Pseudo-colored Live cell PWS image of Hoechst 33342-stained HeLa cells before and after excitation of the dye with UV light. Transformation of chromatin occurs across the whole nucleus within seconds and no repair is observed even after 15minutes (**B**) Hoechst stained and mock-stained cells before excitation and (**C**) the same mock stained and Hoechst stained cells after UV irradiation. (**D**) Minimal (mock) and significant (Hoechst) γH2A.x antibody accumulation. **E)** Distribution of chromatin transformation after UV excitation for Hoechst and Mock stained cells. **F) & (G)** TEM images of Control and Hoechst stained cells confirming nanoscale fragmentation of the chromatin nano-architecture in fixed cells. All pseudo-colored images scaled between Σ=0.01-0.065. Scale bars are all 15μm. Arrows indicate representative nuclei.

Due to this rapid (<10s) chromatin transformation, we hypothesized that the decrease in the mean nuclear Σ was due to the homogenization of the higher-order chromatin organization caused by DNA fragmentation and the resulting nuclear remodeling. To test this hypothesis, we utilized a γ-H2A.X-Alexa488-conjugated antibody to independently monitor the fragmentation of DNA. In Hoechst-stained cells, we observed a drastic accumulation of the γ-H2A.X antibody, whereas we observed little or no localization in the M-S control nuclei [**Figure 2d**]. Additionally, transmission electron microscopy on Hoechst-stained and M-S cells exposed to UV light showed an increase in micron-scale dense chromatin clumps compared with untreated cells [**Figure 2 f&g**]. As previous work has shown that DNA damage causes local chromatin expansion(33), this confirmed our hypothesis that immediate DNA fragmentation was induced by Hoechst-33342 excitation, a phenomenon that is detectable by live cell PWS in real-time without the need for exogenous labels. Subsequently, we compared live cell PWS with phase contrast microscopy to determine if live cell PWS provides information not detectable by other standard, label-free optical modalities [**Figure 3a&b**]. With phase contrast microscopy, no changes in the cell or nuclear structure were detected after excitation of Hoechst 33342 due to its diffraction-limited resolution [**Figure 3b**]. While electron microscopy cannot be performed on live cells, these experiments demonstrate that photo-excitable molecules disrupt the chromatin nano-architecture, which is uniquely detectable in realtime in live cells by live cell PWS.

Next, we investigated the effects of Hoechst staining on the spatial transformation of chromatin nanoorganization as measured by live cell PWS. In particular, we analyzed the spatial distribution of Σ across the nucleus by calculating the two-dimensional spatial autocorrelation, which measures the change in the pixel-to-pixel variability as a function of distance. An increase in the spatial autocorrelation indicates that the nanoscale structure at one pixel has become similar to its neighboring pixels, while a decrease indicates a more locally heterogeneous structure. The size of these clusters of similar nanoscale structures was significantly decreased between 100nm and 1μm after both the addition of Hoechst and its excitation (n=40 from 3 independent experiments) [**Figure 3c**]. This indicates an increase in the spatial microscopic heterogeneity of nanoscopic heterogeneity of the nuclear nanoscale structure (Σ) after Hoechst addition and excitation. Consequently, we found that Hoechst causes a *global alteration* in the chromatin nanoarchitecture independent of its excitation. Not only does this study demonstrate the ability of live cell PWS to sense the heretofore undetectable in live cells alterations in chromatin structure such as double strand DNA breaks, but it also illustrates some of the limitations of the extrinsic labeling approaches, such as Hoechst: even though they have traditionally been used for live cell imaging, these labels alter chromatin structure and lead to DNA damage, which in turn may lead to a perturbation of cell function.

**Figure 3:**
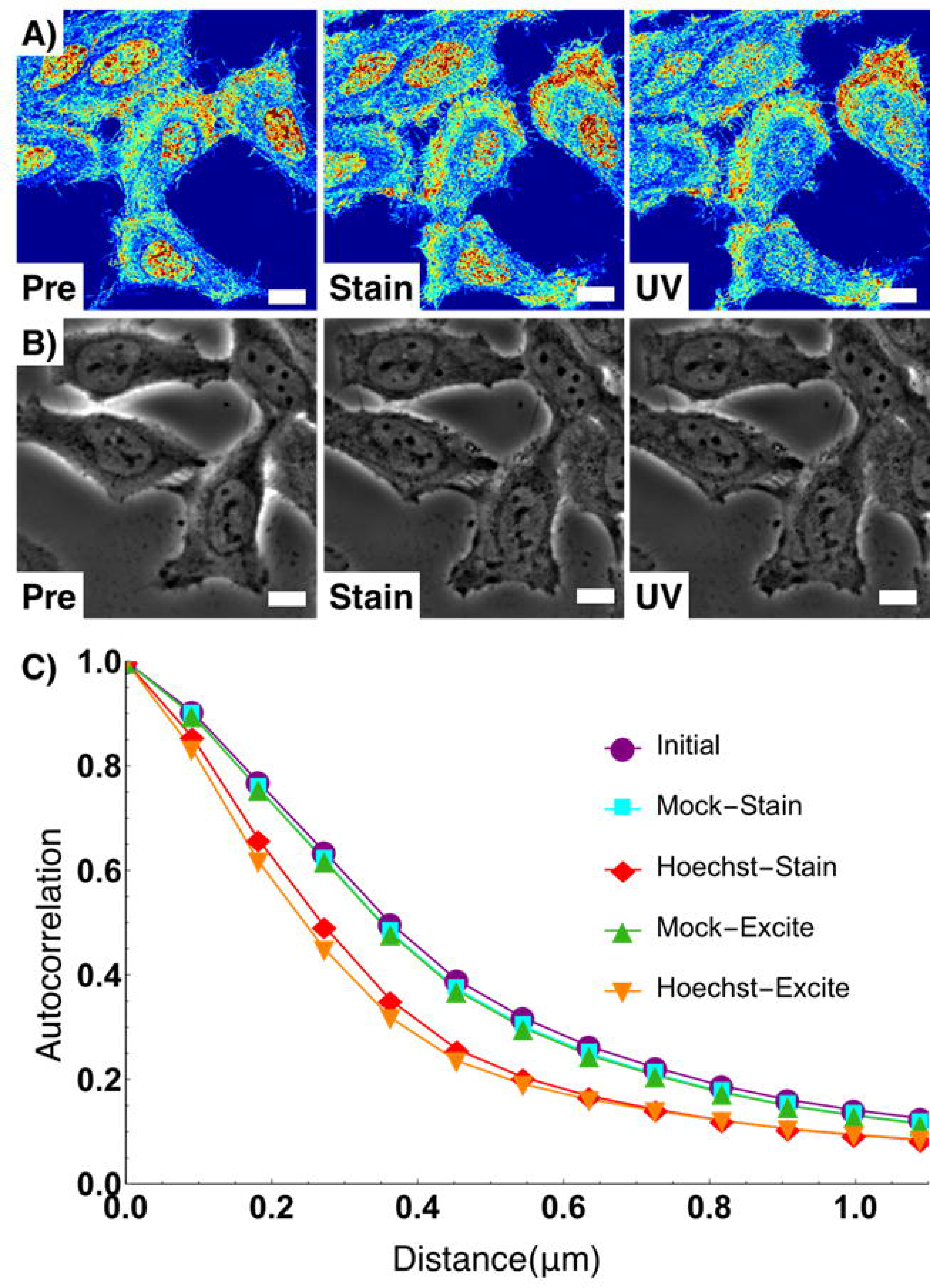
Live cell PWS uniquely detects nano-architectural transformation resulting from Hoechst incubation and excitation. Rows A&B) Live cell PWS (**A**) and Phase Contrast (**B**) cells pre-incubation, 15-minute post-incubation, Hoechst fluorescent image, and after excitation. **C)** Change in the autocorrelation function of Live cell PWS intensity. Hoechst transforms chromatin into a more globally heterogeneous structure. Live cell PWS images are scaled between Σ=0.01-0.065. Scale bars are all 15μm.

Owing to this sensitivity of live cell PWS to detect dynamic changes to the chromatin nano-architecture due to DNA damage, we next applied it to study the temporal dynamics of the cellular nano-architecture under normal growth conditions (**Figure 4a**) in comparison to cells exposed to continuous UV light (**Figure 4b**). UV light is known to cause DNA damage, generate reactive oxygen species, alter receptor-kinase function, and disrupt the cellular membrane. Under normal conditions, chromatin structure can evolve rapidly, with whole-scale changes occurring in minutes (**Figure 4c**, for full videos, see **Supplemental Video 1 & 2**) While the nanoscale topology of chromatin rapidly evolves within any given cell, the organization across the population overall remains stable under normal conditions. In comparison, during continuous UV exposure over 30 minutes, higher-order chromatin structure is degraded after few minutes of exposure (**Figure 4d**, for full video, see **Supplemental Videos 3 & 4**), with pronounced variations in structure over time from cell to cell (with time-lapse measurements performed for 3 independent experiments). As can be observed in **Supplemental Videos 3 and 4**, there are numerous phenomena that occur to the cellular nano-architecture during continuous UV exposure across a distribution of time scales.

**Figure 4:**
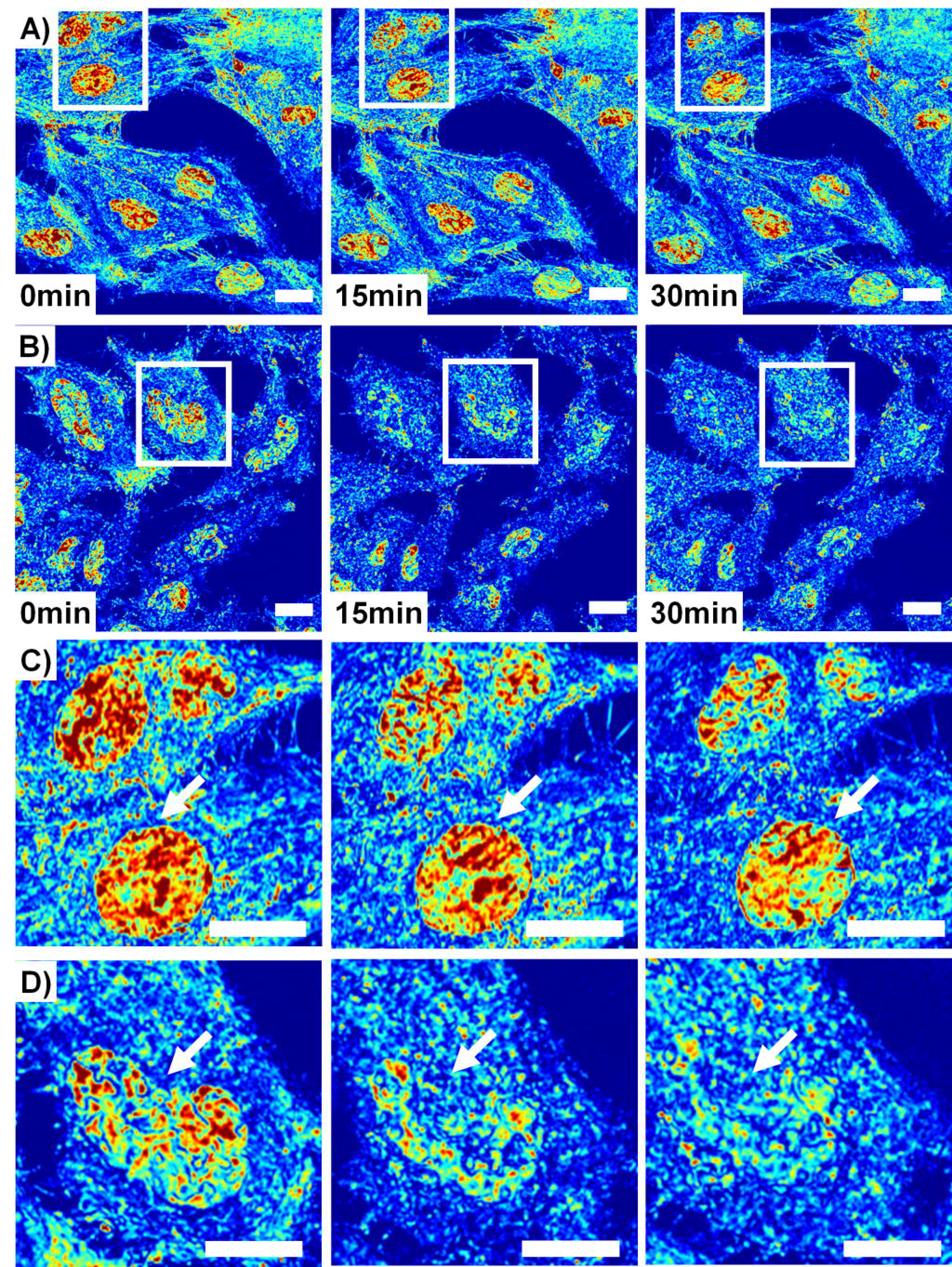
Live cell PWS detects dynamics of nano-architectural transformation under normal and UV-irradiated conditions. Row A) Representative field of view displaying 7 HeLa cells imaged in ~15 seconds using a 63x oil immersion lens, NA=1.4 with Σ scaled to range between 0.01 to 0.065 over 30 minutes of imaging. **Row B)** Representative field of view displaying 7 HeLa cells exposed continuously to UV-light imaged in ~22 seconds using a 63x oil immersion lens, NA=1.4 with Σ scaled to range between 0.01 to 0.065 over 30 minutes of imaging. **Row C)** Inset from field of view in (A) showing the time evolution of two nuclei. Interestingly, chromatin organization is rapidly evolving in time, showing that even at steady state, the underlying structure changes. **Row D)** Inset from field of view in (B) showing the time evolution of one nuclei under UV-illumination. Under UV exposure, homogeneous micron-scale domains form within chromatin, lacking their original higher-order structure. Arrows indicate representative nuclei.

Over the course of 2-3 minutes, there are minimal changes in chromatin and cellular topology due to UV light exposure. However, after approximately ~3 minutes, the chromatin of some cells exposed to UV light undergoes rapid, directional increase in heterogeneity that corresponds with the formation of micron scale homogeneous domains [**Supplemental Videos 3**, **Figure 4e**]. Concurrently, the cytoplasm of the cell is transformed, with cell-cell adhesions retracting and a retreating waveform spreading from the cell periphery toward the nucleus. Finally, a near-instantaneous transition occurs within the cytoplasm, with the changes in the cytoplasmic and chromatin nanostructure spontaneously arresting 20 minutes after exposure [**Figure 4e**]. To capture these temporal dynamics in nanostructure, we performed a kymograph analysis using ImageJ^®^ of a representative cell exposed to UV light in comparison to a control cell. As is shown in **Figure 5a**, over the 30 minutes of exposure to UV, micron-scale homogeneous domains form within the nucleus and the temporal evolution of nanostructure ceases. In comparison, control cells display continuous transformation, with homogeneous and heterogeneous domains transiently forming and dissipating over the time frame of a few minutes [**Figure 5b**]. As is shown in **Figure 5c**, the formation of these large, homogeneous domains that lack high-order structure dominate, resulting in an overall decrease in Σ (average decrease at 30min of 26.9% calculated from 19 nuclei from 3 independent experiments). Interestingly, even under control conditions some cells rapidly demonstrate global changes in their chromatin topology, possibly due to intrinsic molecular variations or due to differential sensitivity to light exposure. Despite these rapid alterations, the overall chromatin structure of the population displays minimal changes over the course of 30 minutes (average 0.2% decrease in Σ from 32 nuclei from 2 independent experiments – additional control experiments with slower acquisition were not included) (**Figure 5c**). Given the multi-modal nature of the current system, these topological variations can be examined and any possible light toxicity further minimized by a variety of well-established methods, including spectral filtration at the illumination source or by using structured illumination.

**Figure 5:**
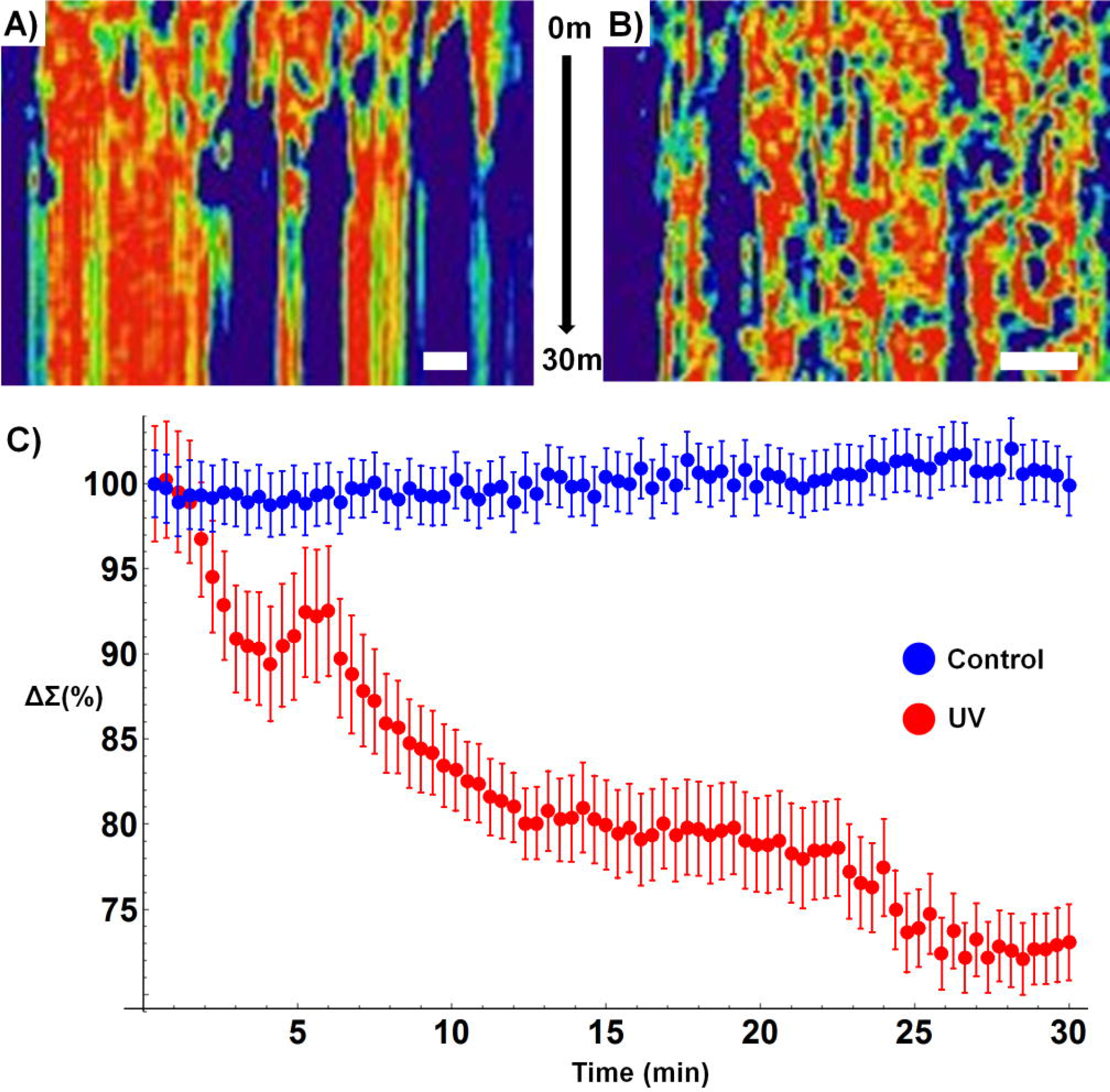
Live cell PWS detects dynamics of nano-architectural transformation under normal and UV-irradiated conditions. A) Kymograph (with the x-axis representing a linear cross-section in x-y plane and the y-axis showing changes over time) representing the temporal evolution of chromatin of a cell exposed to continuous UV-light. Interestingly, nanoscopically homogenous, micron-scale domains form within the nucleus after ~5min of exposure with an overall arrest in structural dynamics. **B)** Kymograph representing the temporal evolution of chromatin of a cell under normal conditions. Under normal conditions, the nanoscale topology of chromatin is highly dynamic, with continuous transitions in structure occurring throughout the nucleus. **C)** Quantitative analysis of nanoscale structure of chromatin of cells under normal conditions (blue, n=32 cells from 2 replicates) and exposed to UV-light (red, n = 19 cells from 3 replicates) for 30minutes. Exposure to UV-light induces overall homogenization of chromatin nanoarchitecture within minutes. Error bars represent standard error. Scale Bar is 5μm.

As a final demonstration of the broad utility of live cell PWS as a tool for studying the complex relationships between cell function and chromatin nano-organization, we studied the effect of alteration of cellular metabolism on higher-order chromatin architecture. The relationship of chromatin structure with mitochondrial function and metabolism has been a major point of focus in recent years. Studies have shown that the cellular metabolic activity is intimately linked to cell replication, tumor formation, DNA damage response, and transcriptional activity(34–37). Therefore, understanding the interplay between the structural organization of chromatin and mitochondrial function is pivotal to understanding numerous diseases.
Recent fluorescence microscopy studies have suggested that impairment of cellular metabolism induces rapid (<15 min) transformation of chromatin(38, 39). However, these studies often require the production of specialized transfection models (H2B-GFP) or the use of DNA-binding dyes such as Hoechst 33342, and as such, are limited in their ability to study multiple cell lines and/or over significant periods of time without perturbing the natural cell behavior(38, 39).

In order to study the link between chromatin structure and mitochondrial function, we employed the protonophore, Carbonyl cyanide m-chlrophenyl hydrazine (CCCP), which is widely used for studies of mitochondrial function due to its disruption of mitochondrial membrane potential (ΔΨ_m_). To explore the role of ΔΨ_m_ reduction on the immediate transformation of the chromatin nano-architecture in live cells, we used two cell lines, HeLa and CHO. Following addition of 10μM CCCP, HeLa cells rapidly lost ΔΨ_m_ whereas CHO cells displayed no significant change as gauged by TMRE fluorescence [**Figure 6a**]. Interestingly, after 15 minutes of treatment with CCCP, we found that addition of 10 μM CCCP produced rapid transformation of chromatin structure in HeLa cells but not in CHO cells [**Figure 6b**]. Critically, in HeLa cells, we observed a decrease in nuclear Σ suggesting homogenization and decompaction in the chromatin structure. Conversely, in CHO cells, we observed no statistical change in chromatin compaction and heterogeneity [**Figure 6c**]. Quantitatively, HeLa nuclei showed a 10% decrease in mean-nuclear E after CCCP (p-value <0.001, n = 31 from 6 independent experiments), whereas the CHO cells displayed no significant increase in mean-nuclear Σ (n=159 cells from 5 independent experiments) [**Figure 6d**]. This transformation suggests that the depletion of mitochondrial membrane potential induces rapid decompaction and homogenization of chromatin nanostructure. Disruption of the ΔΨ_m_ has numerous effects, including the inhibition of mitochondrial ATP synthesis, changes in the production of reactive oxygen species (ROS), altered signal transduction, as well as modification of other mitochondrially produced metabolites (i.e. acetyl and methyl transfer groups). While previous groups have shown that Ψ_m_ is an important determinant of cellular proliferation, to date it has not been shown that loss of Ψ_m_ has a rapid effect on the global chromatin structure. These results show for the first time that the change in Ψ_m_ rapidly regulates the nanoscale organization of chromatin, possibly resulting in the observed decreased proliferative potential of cells over time.

**Figure 6:**
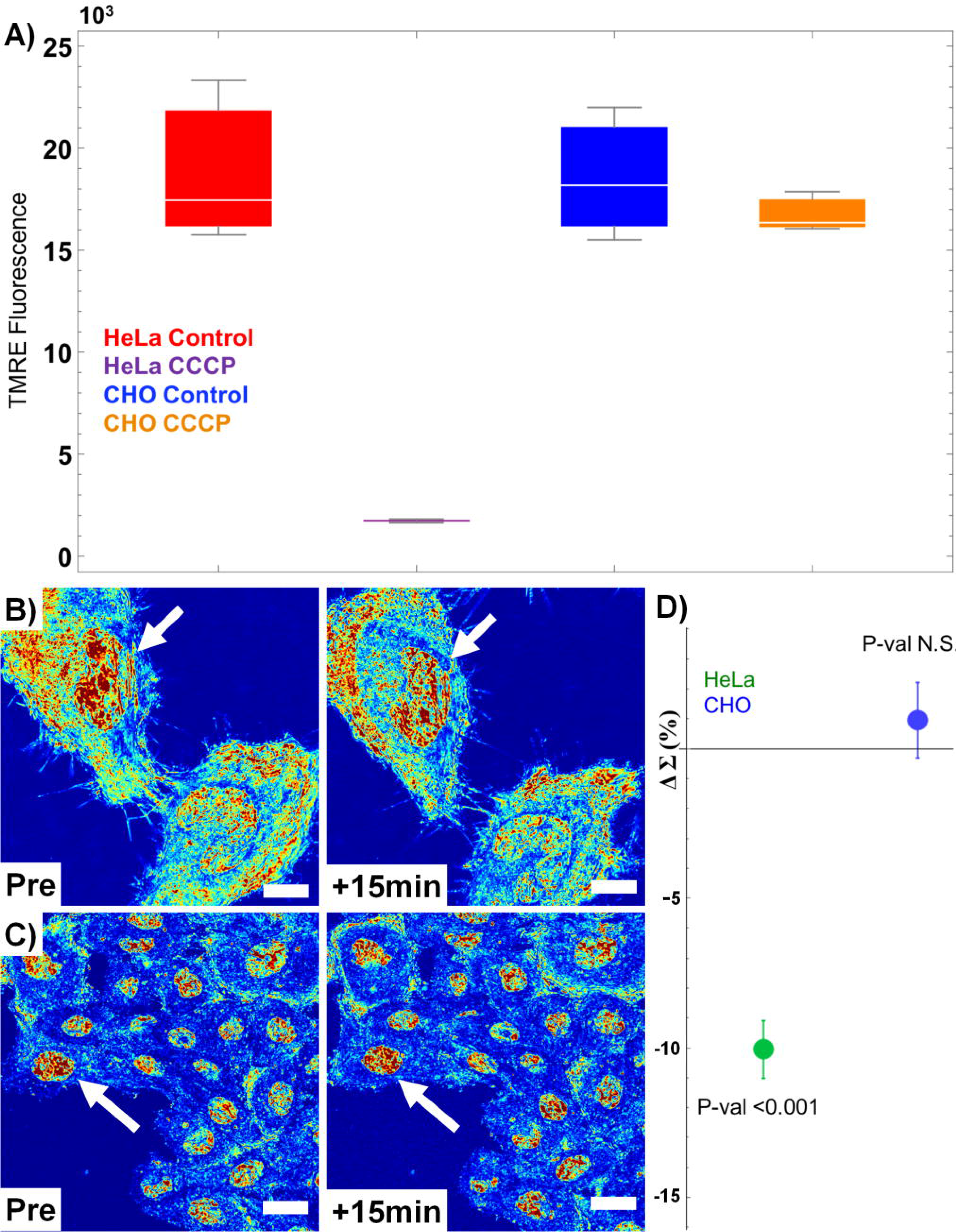
Mitochondrial membrane potential (ΔΨ_m_) is a direct, rapid regulator of chromatin compaction. A) Flow Cytometry showing a 10-fold decrease in Hela cell TMRE fluorescence after 10μM CCCP treatment (p<0.015) and no significant change in CHO cell fluorescence. **Row B)** HeLa and **C)** CHO cells before and 15 minutes after CCCP treatment. **D)** Quantification of the nuclear nano-architecture change in HeLa and CHO cells before and after treatment (HeLa=31cells, 6 replicates and CHO=159 cells, 5 replicates) with standard error bars. Depletion of ΔΨ_m_ induces decompaction and homogenization of HeLa but not CHO chromatin. Live cell PWS images are scaled between Σ=0.01-0.065. Scale bars are all 15μm, arrows indicate nuclei. Arrows indicate representative nuclei.

## Conclusion

In summary, we have extended the application of Partial Wave Spectroscopic microscopy to the study of temporal dynamics of the cellular nano-architecture. Using this technique, we can rapidly quantify the nano-molecular organization in live eukaryotic cells without the use of exogenous labels. While live cell PWS alone is not molecularly specific, it is easily integrated with existing fluorescent methods, providing information that cannot be visualized by existing optical approaches. Furthermore, live cell PWS demonstrates that the nanoscale structure of chromatin evolves rapidly with time, which would significantly transform our understanding of the structure-function relationship between critical processes and chromatin structure, including DNA-repair, replication, and transcription. With this technique, we determined that live cell DNA binding dyes, such as Hoechst 33342, cause rapid destruction of the higher-order chromatin structure at time-scales (seconds) not previously recognized. Paradoxically, this dye is ubiquitously used for the study of cell viability and the presence of DNA damage(40). As a result, live cell PWS is a powerful tool for studying DNA damage/repair and potentially, chemotherapeutic efficacy in live cells. To demonstrate its potential for use in this regard, we studied the temporal dynamics of chromatin during continuous to UV light exposure, showing a transformation in both the temporal and physical properties of the chromatin nano-architecture during UV induced stress.

Additionally, we showed that live cell PWS allows for previously limited exploration of the factors affecting the chromatin nano-architecture by demonstrating differential responses in chromatin structure that depend on the mitochondrial membrane potential. In particular, this illustrates that mitochondrial function is intimately related to chromatin structure in real-time and that live cell PWS can for the first time act as a tool to further investigate the mechanisms of chromatin-metabolic interactions. Live cell PWS is a natural supplement to super-resolution fluorescence techniques, providing quantifiable information about unstained cellular organization to examine the role of the nano-architecture on molecular interactions in live cells. In the future, we envision that live cell PWS can be applied to a broad range of critical studies of structure-function in live cells - leveraging the multimodal potential in conjunction with existing SRM to study: (1) the interaction between chromatin structure and mRNA transport; (2) the accessibility of euchromatin and heterochromatin to transcription factors(41–43); (3) the relationship between chromatin looping, as measured by techniques such as Hi-C, to the physical chromatin structure(6,12,44); (4) why and how high-order chromatin structure changes in cancer(45); (5) the role of nuclear architecture as an epigenetic regulator of gene expression(6, 12, 44); (6) the effect of metabolism on chromatin structure(36, 46); and (7) the role of chromatin dynamics in stem cell development(47, 48).

## Materials and Methods

### Live cell Partial Wave Spectroscopic Imaging

Prior to imaging, media within the petri dishes was exchanged with fresh, RPMI-1640 Media (lacking phenol red pH indicator, purchased from Life Technologies) supplemented with 10% FBS (Sigma Aldrich, St. Louis Missouri). For DNA fragmentation experiments, live cell PWS images were acquired at room temperature (22°C) and in trace CO_2_ (open air) conditions for cells subsequently stained with Hoechst 33342. During acquisition of any time series data (UV and Controls, metabolic perturbation), cells were maintained at physiological conditions for the duration of the experiment. For imaging, a reference scattering spectra was obtained from an open surface of the substrate coverslip immersed in media prior to any cellular imaging to normalize the intensity of light scattered for each wavelength at each pixel. We define Σ as the spectral standard deviation of our measured reflectance intensity normalized to this reference scattering spectra from the substrate-media interface. For Phase Contrast imaging, cells were grown and maintained in the same conditions as cells used for live cell PWS, but images were acquired with a 40x air objective and a transmission illumination beam. Likewise, for wide-field fluorescent imaging, cells were grown in the same conditions but pre-incubated with Hoechst 33342 for 15 minutes prior to imaging. To study the effects of UV illumination on cellular structure and function, cells were continuously exposed to UV light produced from an Xcite 120 LED light source (Excelitas, Waltham, Massachusetts) by removing the 500nm long-pass filter from the illumination path (measurements were performed in triplicate, n = 19). For Hoechst induced DNA damage experiments, significance was determined using Student’s T-test with unpaired, unequal variance on nuclear Δ(Σ) between the conditions indicated in the experiment in both Mathematica v.10 and Microsoft Excel (Microsoft, Redmond, Washington) with n=146 for Hoechst stained HeLa cells from 11 replicates and n=87 for Hoechst stained CHO cells from 5 replicates. For mitochondrial membrane depletion experiments, significance was determined using a two-tailed, paired Student’s T-test on nuclear Σ before and after CCCP treatment using Microsoft Excel (Microsoft, Redmond, Washington) with n = 31 for HeLa cells from 6 independent experiments and n=159 for CHO cells from 5 independent experiments. Each experiment consists of 1-10 independent fields of view for analysis. Sequences of pseudo-colored live cell PWS images were merged into movies using ImageJ. All pseudo-colored live cell PWS images were produced using Matlab^®^ v. 2015b using the Jet color scheme with the ranges indicated in the figure legend. All cells were purchased from ATCC (Manassas, Virginia) unless otherwise noted and imaged in their cell appropriate media supplemented with 10% FBS. Human Umbilical Vein Endothelial Cells (HUVEC) were purchased from Lonza (Walkersville, Maryland) and grown under cell appropriate media formulation on poly-l-lysine coated glass imaging dishes.

### Co-localization

Fluorescence co-localization of organelle specific stains with live cell PWS imaging was performed through manual image alignment of mean-reflectance images produced by live cell PWS acquisition of unstained cells to the cells at the time of acquisition. Background intensity was removed using ImageJ^®^ with using a rolling average of 50 pixels for nuclei and 75 pixels for mitochondria. Threshold intensities for the aligned fluorescence images were then calculated by FindThreshold function in Mathematica^®^ version 10 utilizing Otsu’s algorithm. Co-localized images were produced by the binary mapping of fluorescent images for each stain, pseudo-colored, and scaled by the live cell PWS Σ intensity.

### H2A.X phosphorylation

Co-registration of live cell PWS imaging and DNA strand damage using phospho-histone H2A.X was performed by immunofluorescent staining on three independent experiments. Cells were fixed for 20 min with 4% paraformaldehyde at room temperature, washed twice with phosphate buffered saline and a permeabilization/blocking step was performed with 0.1% Triton X-100 in 1% bovine serum albumin (Sigma Aldrich, St. Louis Missouri) for 20 min. Cells were again washed twice with PBS and then incubated with AlexaFluor 488 conjugated to anti-γH2A.X (serine 139 residue) rabbit monoclonal antibody (Cell Signaling, Beverly, CA) for 30 minutes. Following incubation with the antibody, cells were imaged using the FITC-EGFP filter on the live cell PWS microscope.

### Mitochondrial Membrane Potential Perturbation

HeLa and CHO cells were grown and prepared for live cell imaging as previously described. Cell measurements for a single field of view were sequentially obtained for 3 minutes prior to treatment with CCCP. HeLa (n = 31 from 6 independent experiments) and CHO (n = 159 from 5 independent experiments) cells were treated with 10μM for 15 minutes and imaged before and after treatment. Mock treated cells were incubated with 0.01% DMSO to account for the effect of DMSO solvent on the cells. No significant changes were observed in the mock treated cells for either cell line. Mitochondrial membrane potential ΔΨ_m_ was measured by flow cytometry (BD LSRII at the Northwestern Flow Cytometry Core) for tetramethylrhodamine (TMRE, purchased from Life Technologies, Carlsbad California) stained cells. In brief, cells were trypsinized and immediately stained with 50nM of TMRE for 30 minutes. Cells were washed twice with PBS after staining and suspended in 1ml of PBS. CCCP treated cells were treated for 15 minutes to replicate conditions during live cell live cell PWS imaging. At least 20,000 cells were selected by forward and side scattering channels for each group, with a double elimination of doublets from the final analysis. Mean TMRE intensity from each replicate population was used for representative comparison between treated and untreated groups.

## Acknowledgements

This material is based upon work supported by National Science Foundation Graduate Research Fellowship under Grant DGE-0824162. This work is also supported by the NIH T32 training grants, T32GM008152 and T32HL076139. Additional support was provided by the National Science Foundation Grants No. CBET-1249311. Flow Cytometry was performed by the Northwestern University Flow Cytometry Facility, which has received support from NCI CA060553. Transmission Electron Microscopy was performed at the Biological Imaging Facility at Northwestern University. MDA-MB-231 cells were provided by the O‘Halloran Lab group at Northwestern University.

---

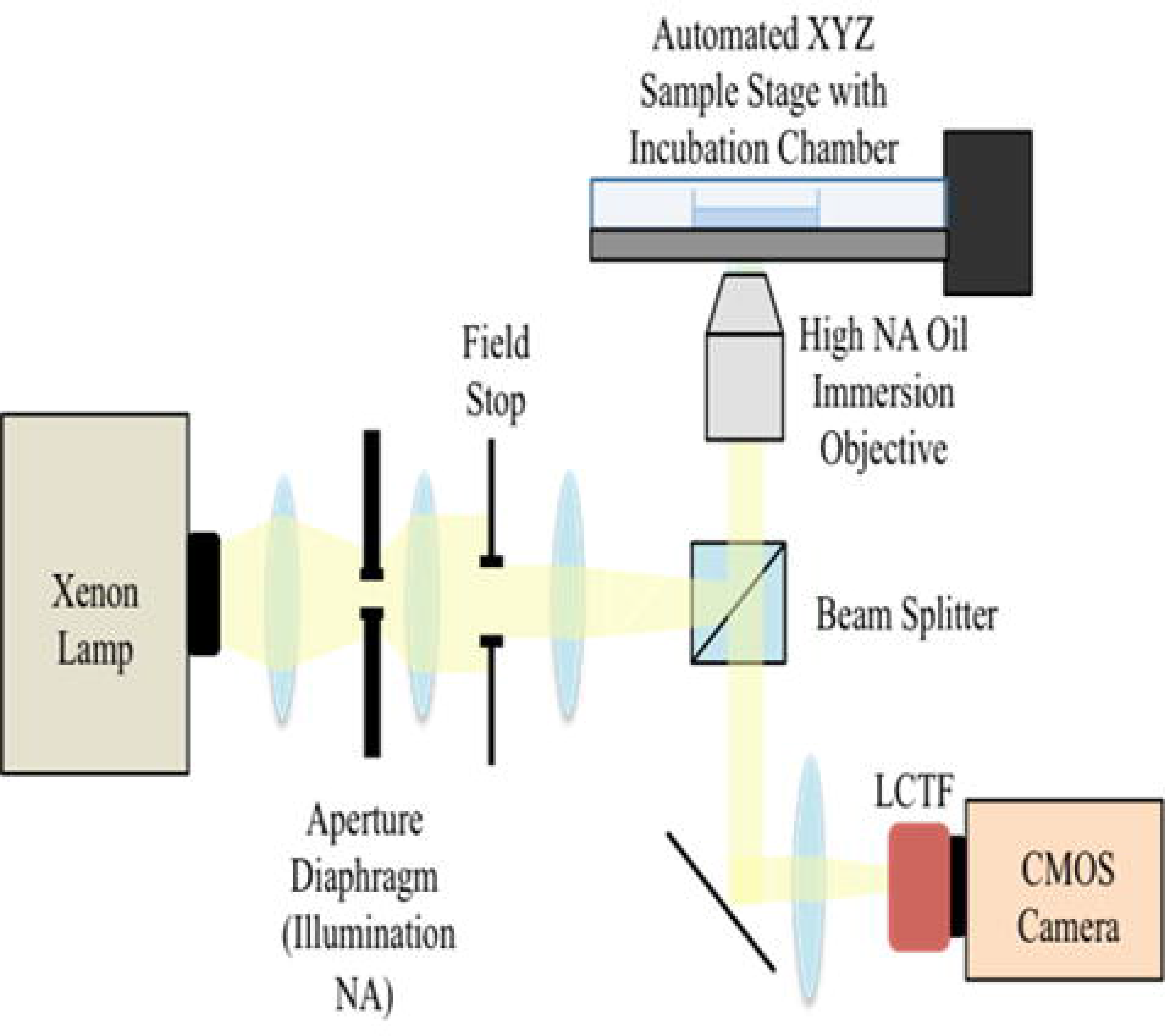

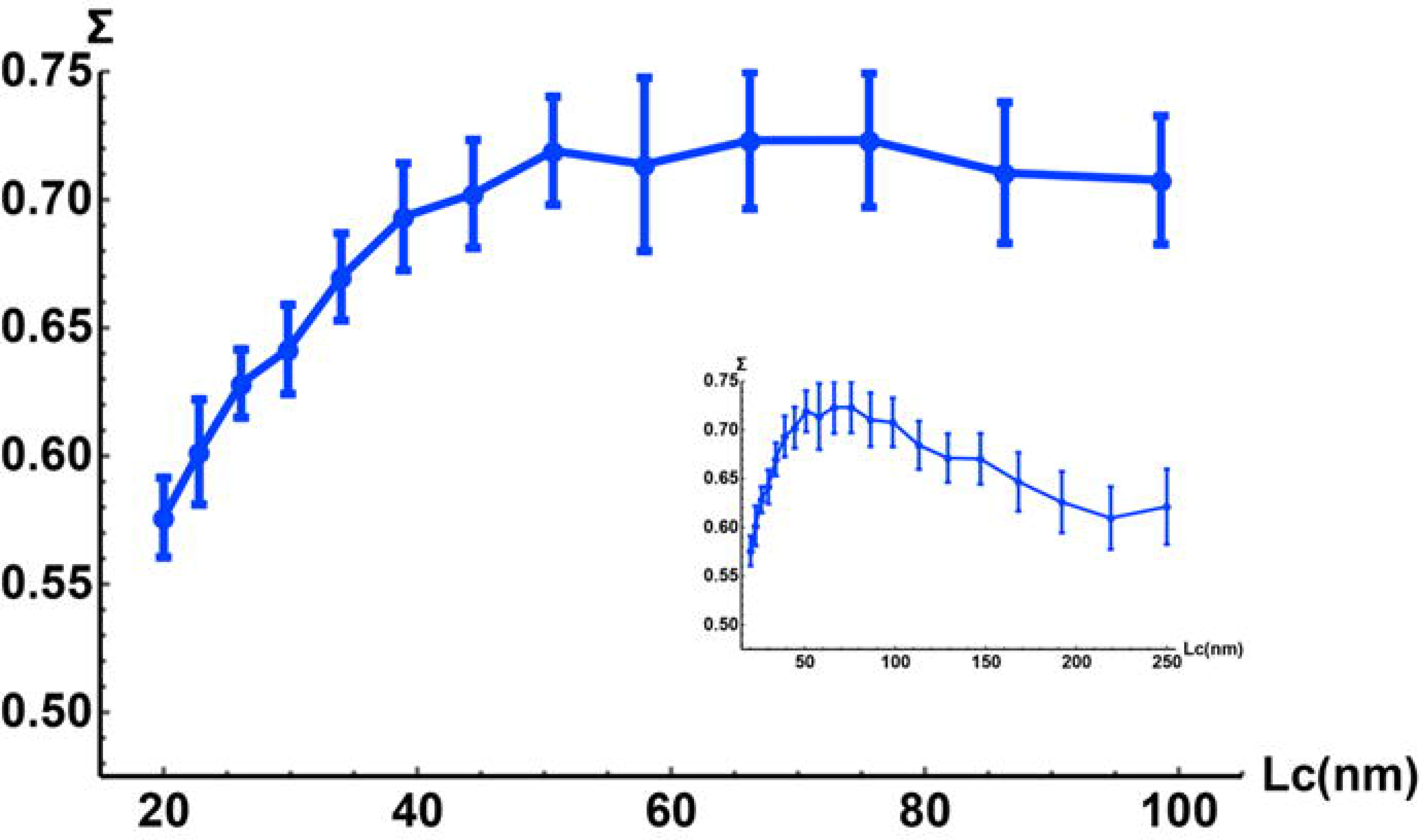

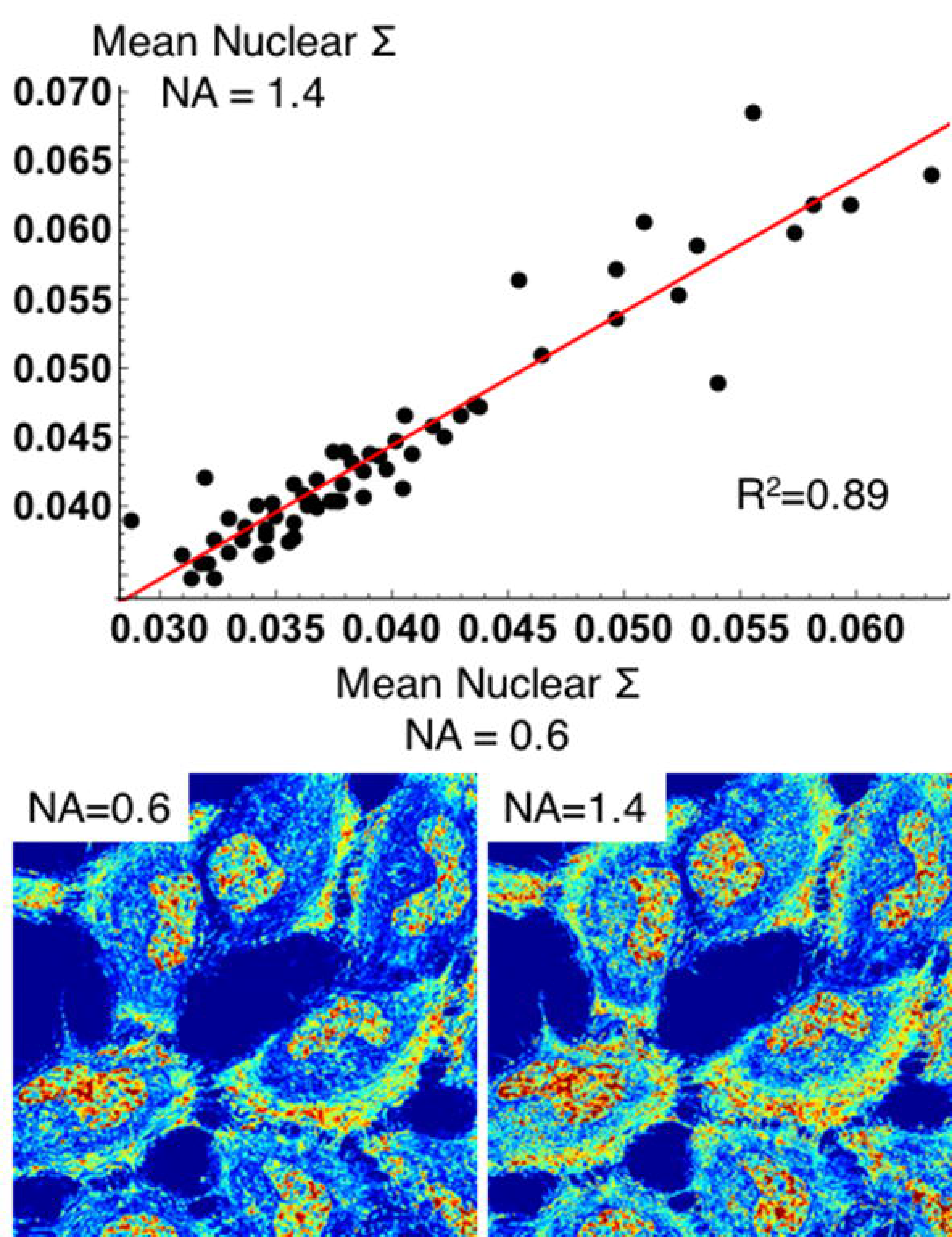

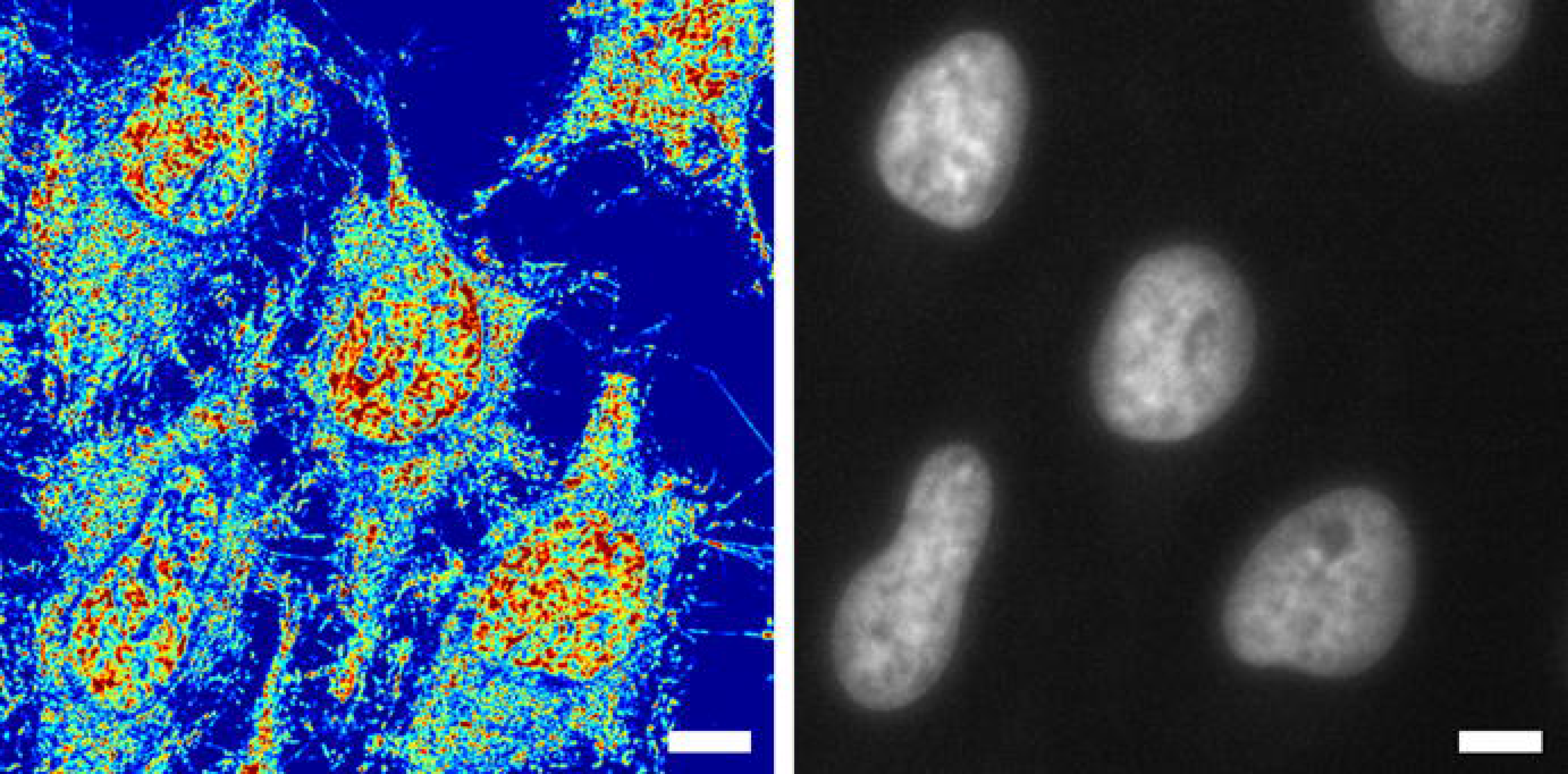

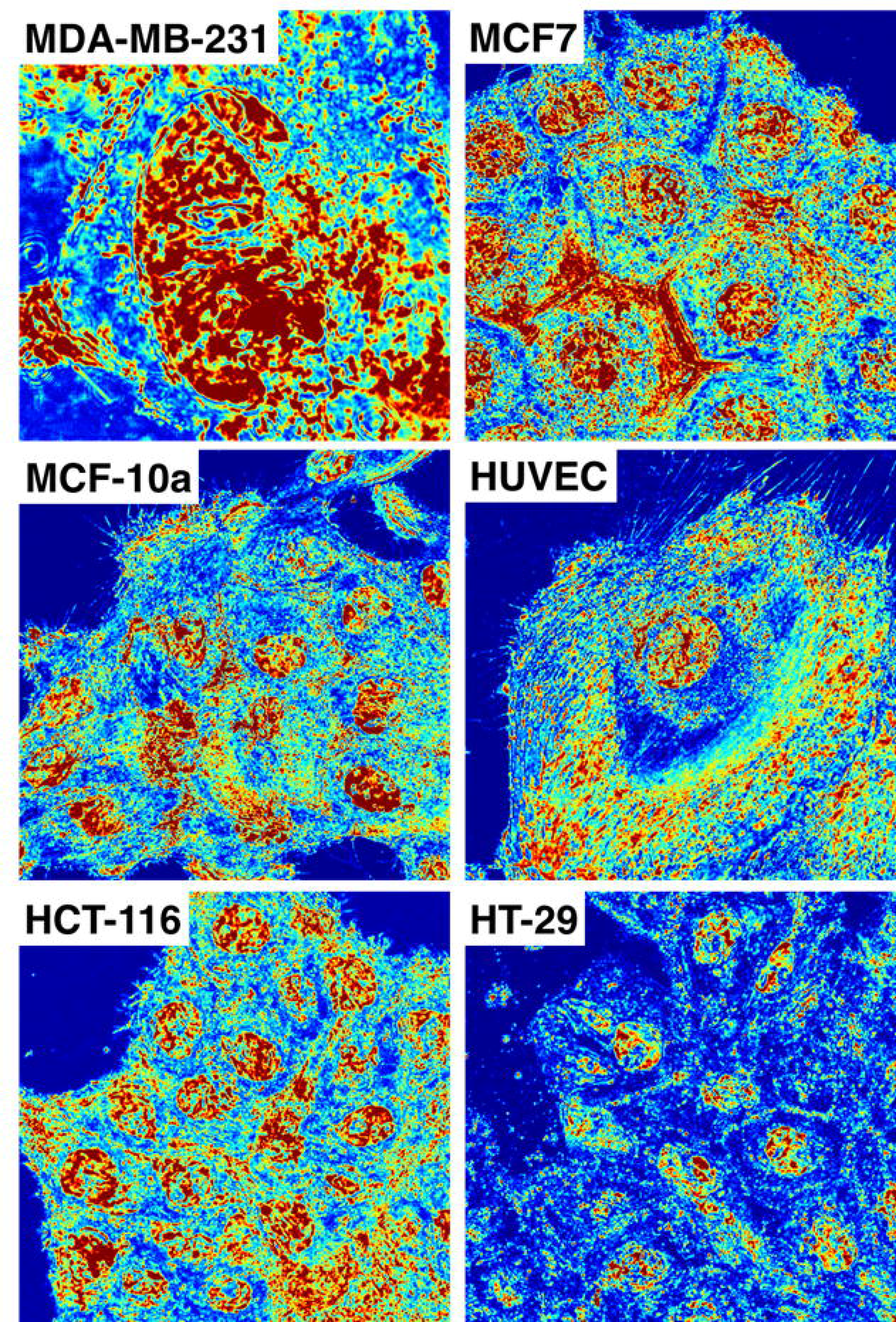

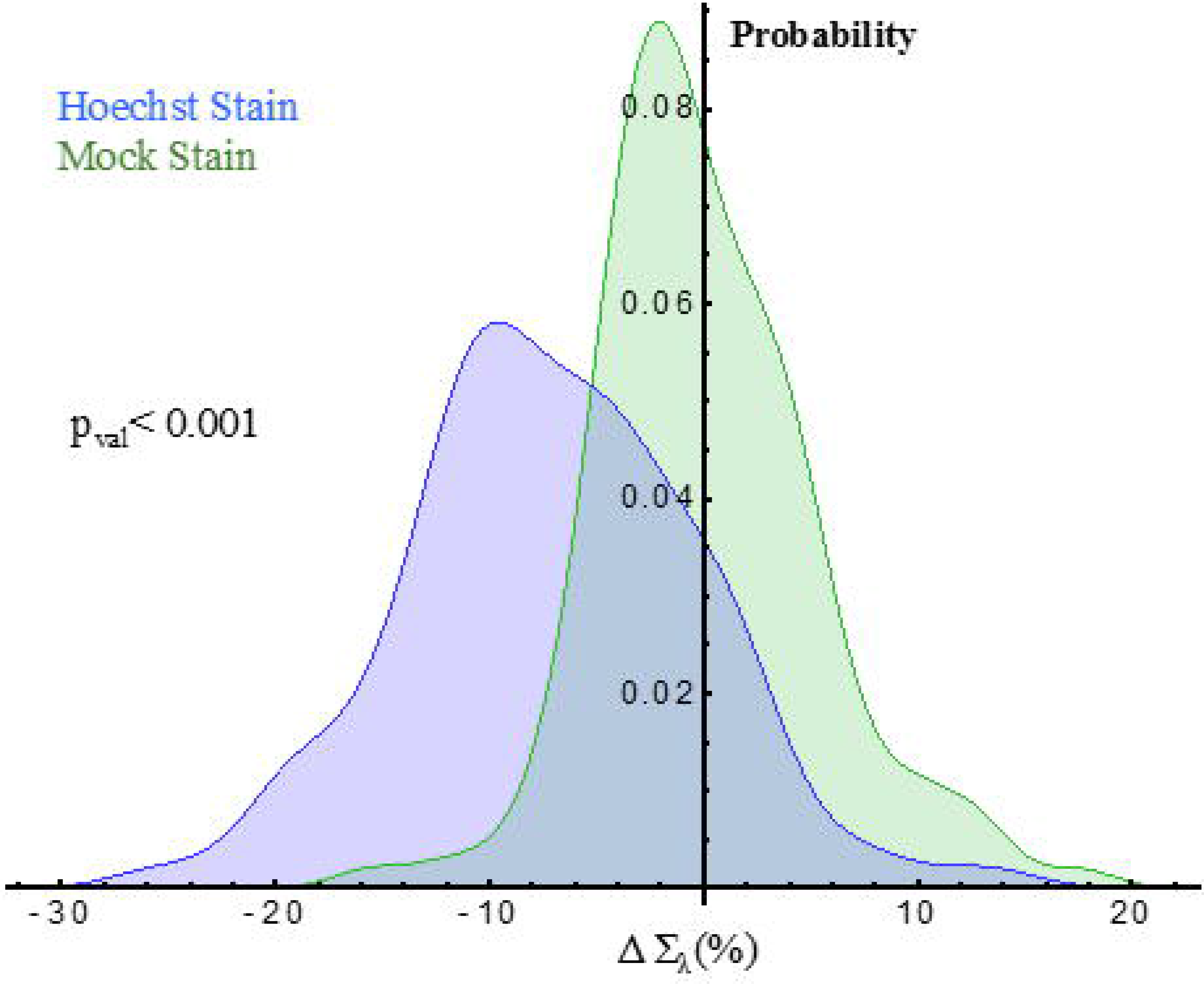

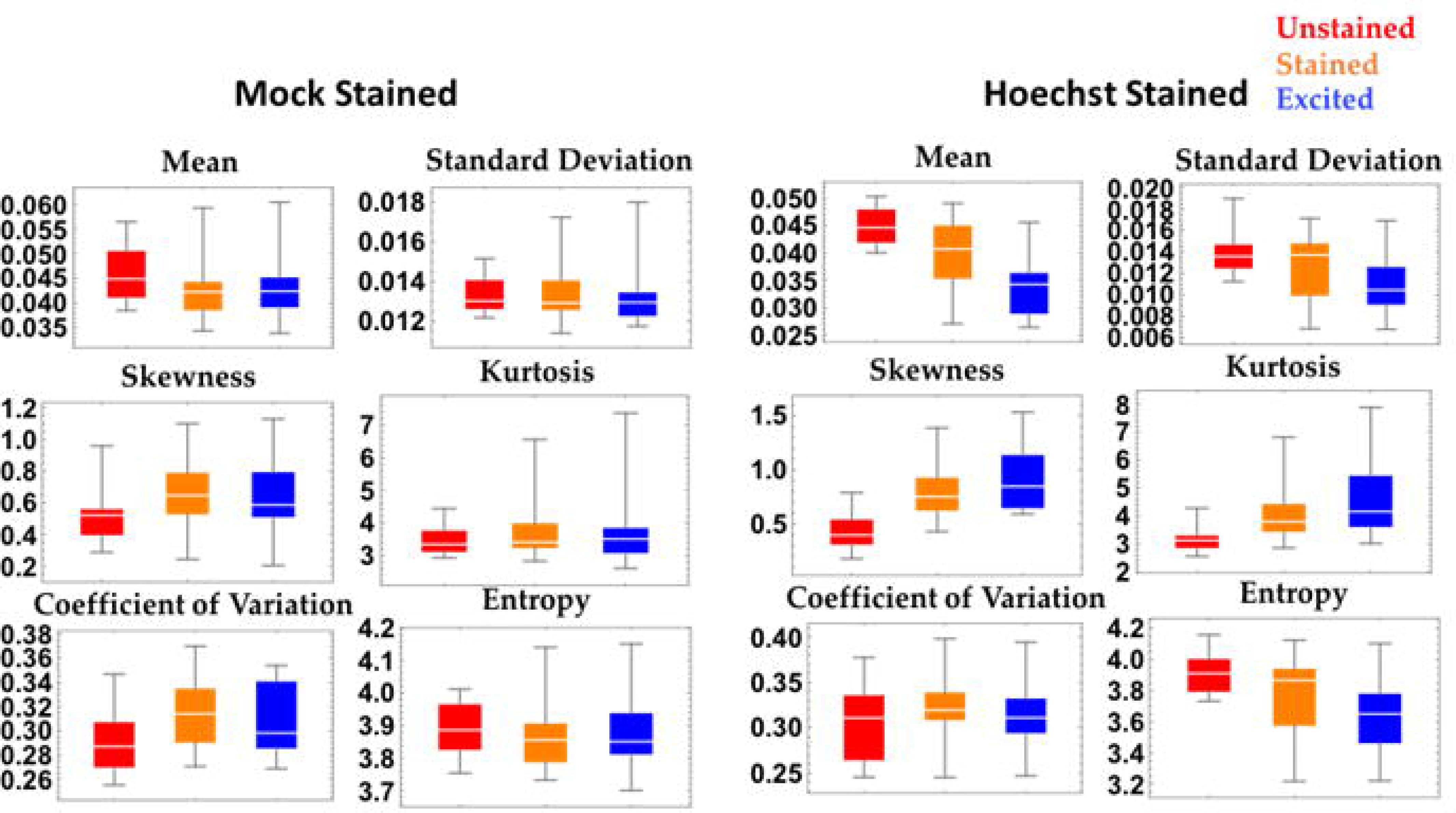

